# Detection of functional protein domains by unbiased genome-wide forward genetic screening

**DOI:** 10.1101/198416

**Authors:** Mareike Herzog, Fabio Puddu, Julia Coates, Nicola Geisler, Josep V Forment, Stephen P. Jackson

## Abstract

Genetic and chemo-genetic interactions have played key roles in elucidating the molecular mechanisms by which certain chemicals perturb cellular functions. Many studies have employed gene knockout collections or gene disruption/depletion strategies to identify routes for evolving resistance to chemical agents. By contrast, searching for point-mutational genetic suppressors that can identify separation- or gain-of-function mutations, has been limited even in simpler, genetically amenable organisms such as yeast, and has not until recently been possible in mammalian cell culture systems. Here, by demonstrating its utility in identifying suppressors of cellular sensitivity to the drugs camptothecin or olaparib, we describe an approach allowing systematic, large-scale detection of spontaneous or chemically-induced suppressor mutations in yeast and in haploid mouse embryonic stem cells in a short timeframe, and with potential applications in essentially any other haploid system. In addition to its utility for molecular biology research, this protocol can be used to identify drug targets and to predict mechanisms leading to drug resistance. Mapping suppressor mutations on the primary sequence or three-dimensional structures of protein suppressor hits provides insights into functionally relevant protein domains, advancing our molecular understanding of protein functions, and potentially helping to improve drug design and applicability.

## INTRODUCTION

In model organisms, genetic screens have long been used to characterize gene functions, to define gene networks, and to identify the mechanism-of-action of drugs (1–4). The genetic relationships identified by such screens have been shown to involve positive and negative feedbacks, backups and cross-talks that would have been extremely difficult to discover using other approaches (5). Currently, the large majority of reported screens in model organisms and in mammalian-cell systems have used gene-deletion libraries and/or methodologies to inactivate gene functions, such as short-interfering RNA, CRISPR-Cas9 or transposon-mediated mutagenesis (6, 7). While powerful, such approaches usually identify loss-of-function phenotypes, and only rarely uncover separation-of-function or gain-of-function mutations. This limitation is significant because such separation- or gain-of-function mutations - which can arise spontaneously or via the action of genotoxic agents - can dramatically affect cell functions or cellular response to chemicals, and can have profound impacts on human health and disease (8, 9). Suppressor screens, either based on lethal genetic deficiencies and/or the use of drugs, have also facilitated the characterization of functionally relevant protein domains and sites of post-translational protein modification through the identification of relevant single nucleotide DNA variants (SNV)s (10).

In their simplest experimental setup, suppressor screens based on point-mutagenesis rely on four tools: (i) a genetically amenable organism or cell; (ii) a selectable phenotype; (iii) a method to create a library of mutants; and (iv) a method to identify mutations driving the suppressor phenotype amongst all the mutations in the library. Reflecting their relative amenability, these screens have mostly been carried out in microorganisms, either bacteria or yeasts, both of which benefit from the ability to survive in a stable haploid state. Despite not being strictly essential for such studies, a haploid state greatly improves the chances of identifying loss-of-function or separation-of-function recessive alleles, which would be masked in a diploid cell state (11). While the first three tools mentioned above are often amenable to a researcher, the lack of fast and efficient methods to bridge the knowledge-gap between phenotype and genotype has discouraged the widespread implementation of suppressor screens based on point-mutagenesis. Indeed, until recently, recessive suppressor alleles could only be identified by labor-intensive methods involving genetic mapping and cloning in yeast, whereas the natural diploid state of mammalian cells largely precluded straightforward SNV suppressor screens in such systems.

Here, we describe an approach to overcome the above limitations that is based on sequencing genomic DNA extracted from various independent suppressor clones, followed by bioinformatic analysis. With small adaptations, this method can be applied to both the budding yeast *Saccharomyces cerevisiae* and other haploid model organisms, as well as to haploid mammalian cells (Figure 1). To highlight the utility of this approach, we describe its application to study resistance to the anti-cancer drugs camptothecin or olaparib, leading to the identification of various mutations in yeast *TOP1* and in mouse *Parp1*, respectively. Importantly, we establish that drug target identification and mechanisms of drug resistance can be unveiled without *a priori* knowledge of the drug target. Furthermore, if a sufficient number of suppressors is screened, this method also allows identification of functional protein domains required to drive drug sensitivity and resistance.

## MATERIALS AND METHODS

### Yeast suppressors of camptothecin sensitivity

*S. cerevisiae* strains used were derived from W303. All gene deletions were introduced by using one-step gene disruption, and were confirmed by PCR and whole-genome sequencing. Full genotypes of strains are described in Supplementary Table 1. Standard growth conditions (1% yeast extract, 2% peptone, 2% glucose, 40 mg/l adenine) were used. Strains YFP1001 and YFP1073 were mutagenized by adding 4.5% ethyl methane sulfonate (EMS) to liquid cultures in logarithmic growth-phase, pelleted by centrifugation and then resuspended in 50 mM K-phosphate buffer for 10 minutes, followed by EMS inactivation with 1 volume of 10% sodium thiosulfate. Suppressors were obtained by plating each strain on 10 YPD plates supplemented with 5 μg/ml of camptothecin (approximately 10^7^ cells per plate). Resistant colonies were picked after 2-3 days of growth at 30°C and isolated by streaking on YPD plates. Suppression was confirmed by retesting camptothecin sensitivity of the isolated strains. Confirmed suppressors were processed for DNA extraction shortly thereafter, in parallel with 2-3 colonies of the initial strain (Figure 1A).

**Figure 1.**
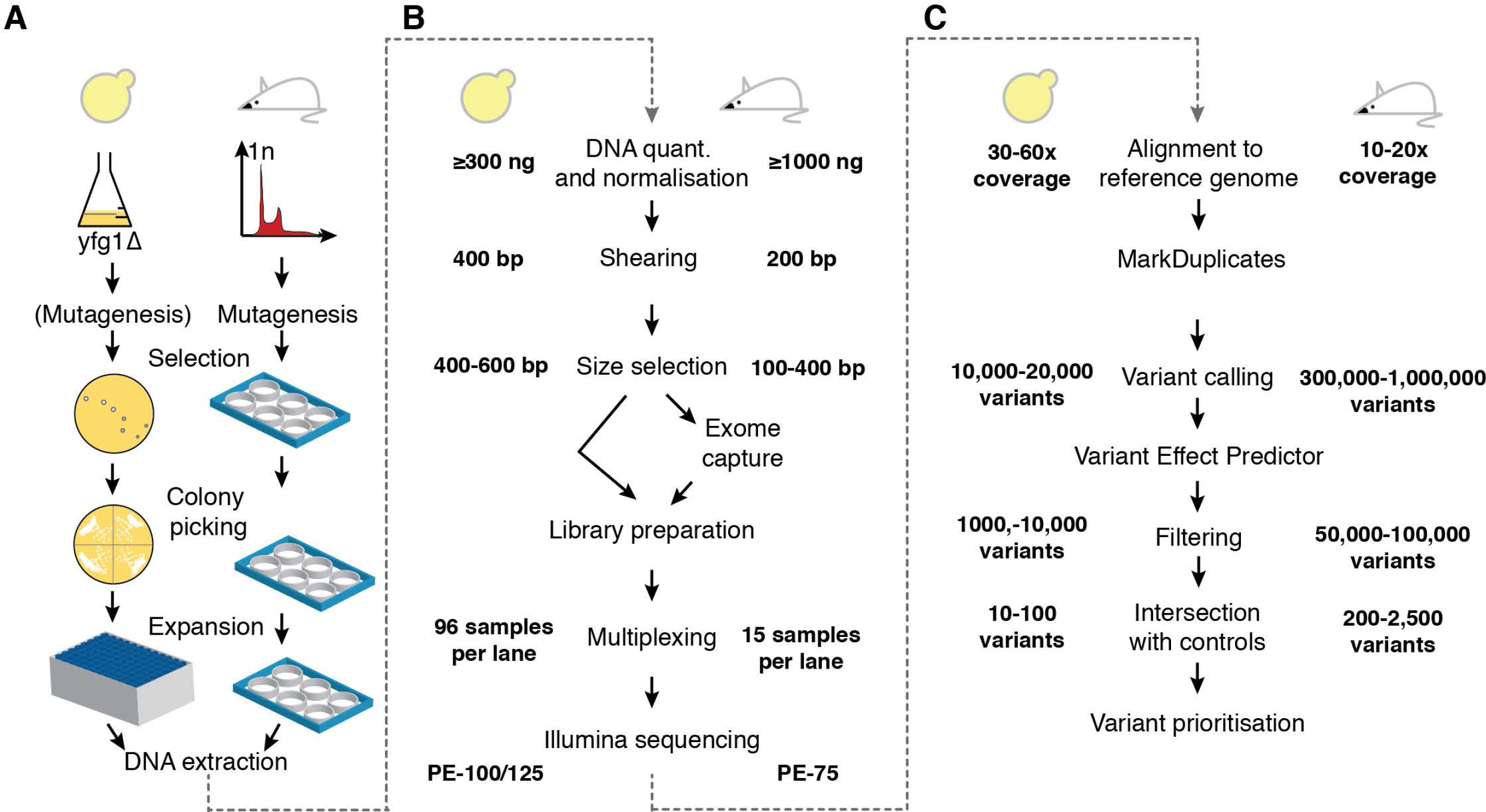
Experimental workflow for a suppressor screen. The typical workflow of a suppressor screen using *S. cerevisiae* (left) or mouse embryonic stem cells (right) is depicted. Details of differences between the two systems are illustrated where appropriate. Variation in mutation numbers for an organism can be due to the choice of background strain, mutagenizing agent and other experimental factors.

### Mouse embryonic stem cell suppressors of olaparib sensitivity

Haploid mouse AN3-12 embryonic stem cells (mESCs) (12, 13) were used for all the experiments and were free from mycoplasma. Cells were grown in DMEM high glucose (Sigma) supplemented with glutamine, fetal bovine serum, streptomycin, penicillin, non-essential amino acids, sodium pyruvate, 2-mercaptoethanol and Leukemia inhibitory factor (LIF). All plates and flasks were gelatinized before cell seeding.

Cell sorting for DNA content was performed on mESCs by using a MoFlo flow sorter (Beckman Coulter) after staining with 15 μg/ml Hoechst 33342 (Invitrogen). The 1n peak was purified to enrich for haploid mESCs.

Mutagenesis with EMS was performed as described previously (14) with the following adjustments: after cell sorting, haploid-enriched cells were grown in DMEM plus LIF for overnight EMS treatment. After EMS treatment, cells were cultured for five passages in DMEM plus LIF and plated into 6-well plates at a density of 5 × 10^5^ cells per well. Cells were then treated with 6 μM of olaparib (AZD2281; Stratech Scientific Ltd.) for 6 days, supplying new medium with olaparib daily. Cells were then grown for another four days without olaparib until mESC colonies could be isolated.

### Genomic DNA isolation

#### *S. cerevisiae* DNA isolation

Resistant colonies were inoculated in 1.8 ml of YPAD in 96-deep-well plates and grown for 48 hours. Pelleted cells were re-suspended in 500μl of spheroplasting solution (1M sorbitol, 0.1M EDTA, 14mM 2-mercaptoethanol, 1mg/ml RNAse A, containing 5 mg/ml zymolyase) and incubated for 2 hours at 37°C. Spheroplasts were subsequently re-suspended in 200 μl of lysis buffer (80% ATL buffer [QIAGEN #19076], 10% Proteinase K [QIAGEN #19133] and 10% RNAse A (10mg/ml)] and incubated overnight (>16 h) at 56°C. Genomic DNA was extracted from the resulting solution by using the Corbett X-Tractor Gene^™^ Robot with the following buffers: AL [QIAGEN #19075; diluted 50% with ethanol], DXW [QIAGEN #950154], DXF [QIAGEN #950163], and E [QIAGEN #950172].

#### Mouse genomic DNA isolation

mESC clones were grown into 12-well plates. After trypsinising and resuspension in 200μl PBS and 200μl Buffer AL [QIAGEN], a proteinase K [QIAGEN, 20 μl] and RNase [QIAGEN, 0.4mg] digestion step was performed (incubating 10min at 56°C). After adding 200 μ1 96-100% ethanol the solutions were applied to QIAamp Mini spin columns following the QIAamp DNA Blood Mini Kit [QIAGEN] manufacturers protocol from there. Genomic DNA was eluted from the columns using 200μ1 distilled water. A second elution was performed if the yield of the genomic DNA obtained was lower than 2 μg. Genomic DNA was stored at −20°C short-term before sequencing.

### Illumina library preparation and sequencing (Figure 1B)

Extracted DNA was tested for total volume, concentration and total amount by using gel electrophoresis and the Quant-iTTM PicoGreen® dsDNA Assay Kit (ThermoFisher Scientific). Genomic DNA - 500 μg (yeast) or 1-3 μg (mouse) - was fragmented to an average size of 100-400bp (mouse) or 400-600bp (yeast) by using a Covaris E210 or LE220 device (Covaris, Woburn, MA, USA), size-selected and subjected to DNA library creation via established Illumina paired-end protocols. Adaptor-ligated libraries were amplified and indexed via PCR. A portion of each library was used to create an equimolar pool comprising 45 indexed libraries for mouse samples, and 96 indexed libraries for yeast samples. For mouse whole-exome sequencing, pools were hybridized to SureSelect RNA baits (Mouse_all_exon; Agilent Technologies).

Mouse libraries were sequenced at 15 samples per lane. Yeast libraries were sequenced at up to 96 samples per lane. Libraries were sequenced by using the HiSeq 2500 (Illumina) to generate 75 (mouse), or 100/125 (yeast) base paired-end reads according to the manufacturer's recommendations.

### Analysis of DNA sequence data to identify suppressor mutations (Figure 1C)

#### Alignment of DNA sequencing data

Sequencing reads were aligned to the appropriate reference genome using BWA aln (v0.5.9-r16) (15). The *S. cerevisiae* S288c assembly (R64-1-1) from the Saccharomyces Genome Database was obtained from the Ensembl genome browser. For mouse samples, the *Mus musculus* GRCm38 (mm10) was used. Where appropriate, all lanes from the same library were merged into a single BAM file, and PCR duplicates were marked by using Picard Tools (Picard version 1.128). The quality of the sequencing data post-alignment was assessed by using SAMTools stats and samtools flagstats (1.1+htslib-1.1), plot-bamstats, bamcheck and plot-bamcheck(16).

#### Variant calling, consequence annotation and filtering

SNVs and small insertions/deletions (INDELs) were identified using SAMtools mpileup (v.1.3) (16), followed by BCFtools call (v.1.3) (16). The following parameters were used for SAMtools mpileup: -g -t DP,AD -C50 -pm3 -F0.2 -d10000. Parameters for BCFtools call were: -vm -f GQ. All variants were annotated by using the Ensembl Variant Effect Predictor (VEP) v82(17). To exclude low quality calls, variants were filtered by using VCFtools vcf-annotate (v.0.1.12b) (18) with options -H -f +/q = 25/SnpGap=7/d=5, and custom filters were written to exclude variants with a Genotype Quality (GQ) score of less than 10. In the case of whole-exome sequencing data, variants called outside of targeted regions were excluded. INDELs were left-aligned using BCFtools norm (16).

#### Removal of background mutations

Variants that confer resistance are absent in the initial strain/cell line, as it is sensitive to the drug used. Bedtools intersect was therefore used to remove variants present in any *S. cerevisiae* control samples to eliminate variation of the background relative to the reference genome from the dataset. Variant calls from mouse samples were filtered by removing all variants identified in sequencing data of three olaparib-sensitive AN3-12 clones using VCFtools vcf-isec (18); INDELs were further verified by using the microassembly-based variant caller Scalpel (20). To address the high false positive rate in INDEL variant calls, only the INDELs that were identified by both variant callers and have passed the filters were retained.

#### Variant prioritization

Variants were prioritized by their Ensembl VEP (17) predicted consequence: we retained variants predicted to cause a frameshift, a premature stop codon, a missense mutation, a lost start/stop codon, a synonymous mutation, an in-frame insertion or deletion, and in case of mouse data those annotated to affect splice donor/acceptor bases. Genes were prioritized by ranking them by the number of distinct mutations identified in each gene.

Missense mutations identified in genes of interest were ranked by using predictions of PROVEAN/SIFT (21–24) and PredictProtein (25). Scores below −2.5 for PROVEAN, above 50 for PredictProtein and below 0.05 for SIFT indicate likely deleterious consequences to protein function.

### Analysis of olaparib resistant cell lines

#### Molecular modeling

Molecular models were generated by using pymol. Crystal structure data were obtained from RCSB Protein Data Bank (PDB). The codes for PARP1 structures were 4DQY (human PARP1 without Zn2 or BRCT domain) and 3ODC (human PARP1 Zn2).

#### Antibodies for immunoblotting

Anti-PARP1 (Cell Signalling, 9542; 1:1000 in TBST 5% milk), anti-HDAC1 (Abcam ab19845; 1:1000 in TBST 1% BSA), anti-PAR (Trevigen 4336- BPC-100, 1:1000 in PBST[0.05%Tween-20] 5%milk), and anti beta-actin (Cell Signaling #4970, 1:1000 in TBST 5% BSA) were incubated with western-blot membranes at 4°C overnight with gentle rocking.

#### DNA binding assays

DNA binding of PARP1 was assayed by using a 26-bp palindromic DNA duplex (5’GCCTACCGGTTCGCGAACCGGTAGGC3’, (26)) immobilized on Dynabeads^™^ M-280 Streptavidin (Invitrogen, 10mg/ml). Individual incubations used 500 μg of protein extract and a buffer containing 10mM HEPES (p.H. 7.4), MgCl2 (1.5mM), 25% glycerol, KCl (200mM), EDTA (0.2mM), Roche protease inhibitor cocktail (0.7X), DTT (0.5mM) and AEBSF (0.1mM). Experiments were repeated at least twice.

#### PARrylation assay

Cells were treated with 6μM Olaparib overnight, followed 1mM H_2_O_2_ for 10 minutes in the dark, washed with ice-cold PBS and collected. Cells were lysed in Laemmli buffer (120mM TrisHCl pH6.8, 4% SDS, 20% glycerol) and lysates were separated on 4-20% Tris-Glycine gradient gel followed by transfer onto PVDF membrane. Membranes were immunoblotted with the appropriate antibodies. Experiments were repeated at least twice.

## RESULTS

### Identification of *TOP1* mutations conferring camptothecin resistance

To demonstrate the utility of the procedure described above, we sought to identify mutations imparting resistance to camptothecin, a DNA topoisomerase 1 inhibitor (27–29). To do this, we employed yeast strains carrying mutations inactivating pathways required for camptothecin resistance. These specific mutations (*rad50S*, *sae2-F267A*, *rtt107Δ*, *tof1Δ*, *sae2Δmre11-H37R-tel1Δ*) were chosen for their ability to induce camptothecin hypersensitivity (30–33). To increase the variety of potential suppressor mutations, two of the five strains used were mutagenized with ethyl methane sulfonate (EMS), an alkylating agent that induces SNVs (34), before plating them in the presence of camptothecin. In all cases, camptothecin-resistant colonies were readily detectable after 2-3 days of growth at 30°C.

Genomic DNA sequencing of the resistant clones highlighted *TOP1* as the gene carrying the largest number of unique mutations in our dataset, as expected for it being the drug target. The second most mutated gene — *PDR1* —carried 11 unique mutations, 10 of which did not co-occur with mutations in *TOP1*, whereas all the mutations found in the third most mutated gene *(GLT1)* co-occurred with mutations in either *TOP1* or *PDR1* (Figure 2A **and data not shown**). Globally, out of the 251 yeast strains sequenced, 191 contained one or more mutation in *TOP1* (Figure 2B, **light yellow**). Furthermore, by manual inspection, we found that 27 additional strains carried mutations in *TOP1* (Figure 2B, **dark yellow**); the inability to automatically detect these mutations was caused by the fact that these strains were either not pure clones, or they carried large (>25bp) deletions in *TOP1* (Figure 2B **and** Supplementary Figure 1A). To the list of *TOP1*-mutated suppressors strains, we added another 38 suppressors bearing *TOP1* mutations that we had identified in previous, published screens (31, 35), bringing the total number of *TOP1* mutants analyzed to 256.

**Figure 2.**
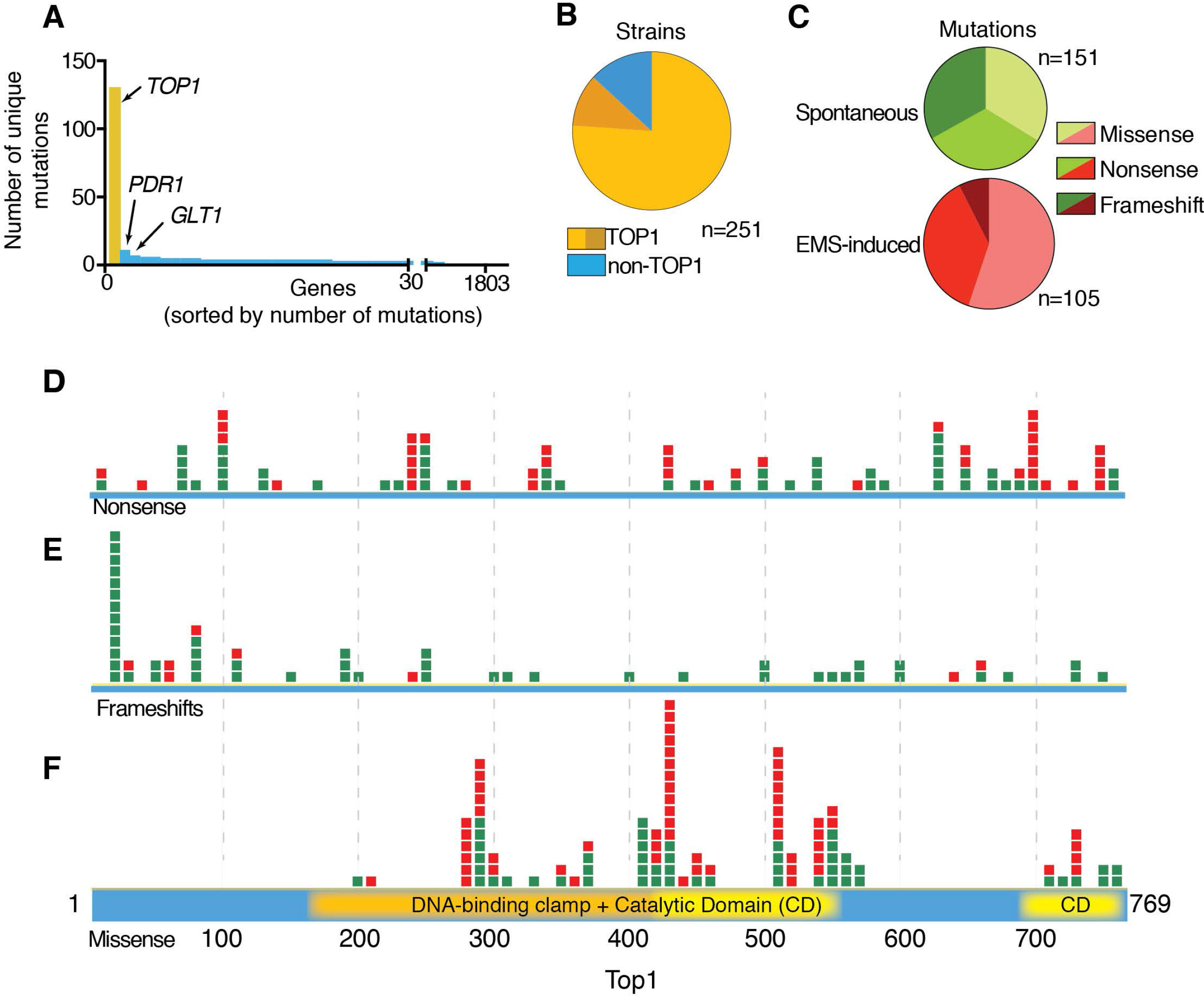
Screening for camptothecin resistance highlights TOP1 and its functional domains. (A) Genes found mutated in screenings for camptothecin resistance sorted by the number of non-synonymous, independent mutations identified in each gene. (B) Fraction of strains identified as mutated in *TOP1* by initial analysis (light yellow) and upon manual inspection of the *TOP1* gene (dark yellow); blue represents clones where no *TOP1* mutation was identified. (C) Mutation types identified in EMS mutagenized samples and non-EMS mutagenised samples. Location of nonsense (D), frameshift (E) and missense (F) mutations in the *TOP1* gene with respect to the primary protein sequence. Mutations identified in EMS-treated samples are colored red, while those from non-mutagenized samples are colored green.

Missense, nonsense and frameshift *TOP1* mutations were roughly equally represented in the non-mutagenized samples. However, where samples had been mutagenized with EMS the vast majority of mutations were nonsense or missense base substitutions (Figure 2C). In the few cases in which the same suppressor clone contained missense and nonsense mutations in *TOP1*, the suppressive effect was attributed to the gained STOP codon.

When the positional distribution of each mutation type was plotted, nonsense and frameshift mutations were shown to be quite evenly distributed along the length of the *TOP1* open reading frame (Figure 2D **and** 2E). The prediction is that such mutations either result in null alleles - as the prematurely-terminated messenger RNA (mRNA) would be degraded by nonsense-mediated decay mechanisms (36) - or would give rise to an unstable protein or a truncated version that could retain partial activity. Since the Y727 residue is essential for the catalytic activity of Top1, truncation before this residue is predicted to produce a non-functional protein (37, 38). As might be expected, the distribution of nonsense mutations loosely correlated with them arising from codons in the open reading frame that only required one nucleotide change to change them to a STOP codon (Supplementary Figure 1B). Notably, the observed enrichment of frameshifts near the 5’ end of the *TOP1* transcript was localized to an 8-nucleotide homopolymeric adenine tract that is presumably particularly susceptible to mutagenesis (Supplementary Figure 1C).

In striking contrast to the situation with nonsense or frameshift mutation, missense mutations were localized to specific regions of the *TOP1* protein-coding sequence, overlapping with known functional domains of Top1. Indeed, the vast majority of mutations identified localized within three distinct regions of the larger DNA binding and catalytic domain, while a minority was located in the smaller C-terminal domain, essential for catalysis (Figure 2F).

Functional consequences of the amino acid residue changes induced by missense mutations were assessed by using PROVEAN and PredicProt (24, 25). These tools use chemical properties of amino acid residues and phylogenetic conservation to predict whether or not a particular substitution is likely to be functionally tolerated by the protein analyzed. Both these methods suggested that the vast majority of the *TOP1* mutations we identified in camptothecin resistant strains were likely to produce deleterious effects (PROVEAN score < −2.5; PredictProtein score >50) (Figure 3A). Notably, missense mutations located in the C-terminal domain of Top1 affected both conserved and non-conserved residues and were primarily positioned in the vicinity of the catalytic residue Y727, although three substitutions were closer to the C-terminus of the protein (Figure 3B).

Top1 binds to DNA via a clamp-like mechanism in which DNA binding stimulates a conformational change in the protein. Thus, opposable “lip” domains encircle the DNA, stabilizing binding through establishing non-covalent protein-DNA and lip-lip interactions (Figure 3C) (39, 40). Approximately two thirds of the missense suppressor mutations identified in the DNA binding domain clustered within the Lip1 and Lip2 regions, highlighting their importance for Top1 function (Figure 3D; the Lip2 domain also contains an active-site residue, R420). Remaining mutations clustered between amino acid residues 500 and 600, which encompass the end of the DNA binding/catalytic domain and the base of the coiled-coil linker domain. In this region two other active site residues (R517 and H558) are located (Figure 3D).

Collectively, these results showed that even with no *a priori* knowledge, our approach for identifying suppressor strains and associated mutations would have identified Top1 as the likely target of camptothecin and would have highlighted the critical Top1 domains functionally relevant for Top1 activity and drug hypersensitivity.

### Identification of mouse *Parp1* mutations conferring olaparib resistance

Based on a similar approach to that described above, we recently identified genes whose mutation in haploid mammalian cells causes resistance to the anti-metabolite drug 6-thioguanine(41). To further highlight the wider applicability of our approach in mammalian cell systems, we carried out a screen to identify mutations that allow haploid mouse cells to survive in the presence of the anti-cancer agent olaparib, a potent small-molecule inhibitor of the DNA-repair protein PARP1 (Poly ADP-ribose polymerase 1) (42, 43). Thus, wild-type, haploid mouse embryonic stem cells (mESCs) were mutagenized by using EMS, and mutant libraries were screened for resistance to olaparib (Figure 1A). Forty-five olaparib-resistant clones were isolated and subjected to whole-exome sequencing.

**Figure 3.**
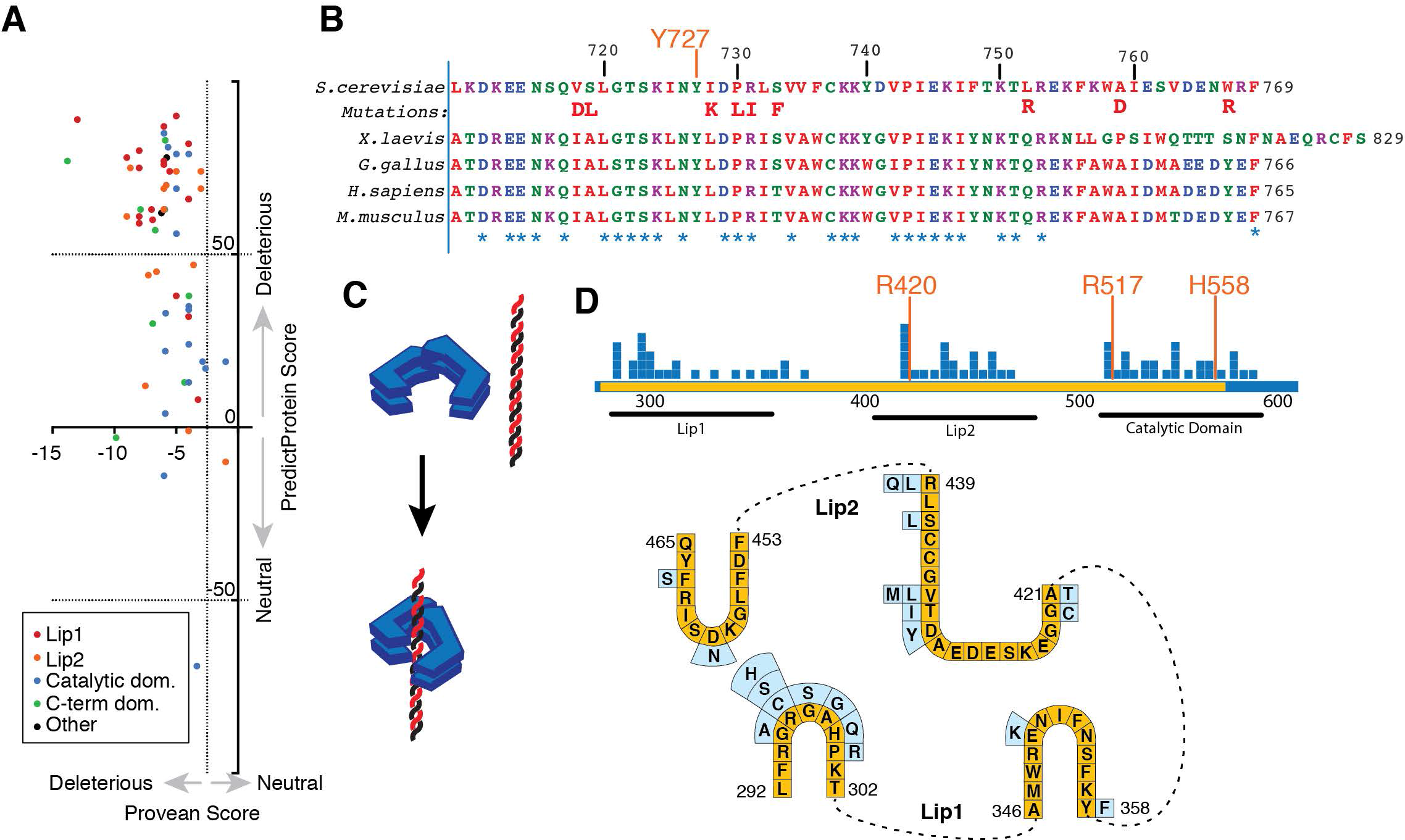
Mutations conferring camptothecin resistance affect key functional residues in *TOP1*. (A) Computational predictions for consequences of *TOP1* missense mutations as predicted by PredictProtein (y-axis; a score above 50 is considered deleterious) and PROVEAN (x-axis; a score below −2.5 is considered deleterious). Datapoints are colored by the domain that the missense mutations occur in: red and orange denote mutations falling within the Lip1 and Lip2 domains, respectively, blue represents the catalytic domain before the Linker, green the C-terminal catalytic domain. (B) Multi-species alignment of the Top1 protein catalytic domain, with the tyrosine residue critical for catalysis (Y727) being highlighted in orange and mutations identified in this screen being highlighted in boldface red. (C) Model of the Top1 mode of DNA binding. The protein wraps around to doublestranded DNA with its two Lip domains, like a grasping hand. (D) Missense mutations identified in the DNA-binding clamp and active site region with respect to the Lip DNA-binding regions and the active site. The region critical for DNA binding and catalysis is highlighted in yellow, and mutations are represented by blue squares. Below is a depiction of the sequence and loop structure of the Lip1 and Lip2 regions in yellow, with specific mutations indicated in light blue.

Analysis of ensuing sequence data for putative, acquired mutations, revealed *Parp1* as the most mutated gene in the dataset with 25 different mutations detected (Figure 4A, **Supplementary Table 3**). Globally, 40 out of the 45 clones harbored *Parp1* mutations (Figure 4B, **Supplementary Table 3**). Further manual examination of the aligned sequencing data from the five remaining clones revealed that four of these also likely carried mutations affecting PARP1 (Supplementary Figure 2). Two of those five (A7, B7) likely carry the R138C missense mutation identified in another clone (Supplementary figure 2A, Figure 4C), while two other clones (A9, H10) likely harbored nonsense mutations at codon 341 (Supplementary Figure 2B). Importantly, mutations in the second and third most mutated genes (*Ttn* and *Plch1* with 9 and 5 different mutations, respectively) never occurred in isolation in the absence of *Parp1* mutations, while *Parp1* mutations also occurred in the absence of *Ttn* or *Plch1* mutations. These data thus highlighted how such analysis would have identified PARP as the likely prime driver of olaparib sensitivity without any knowledge about the drug’s mechanism-of-action (see below for further discussion).

Of the *Parp1* mutations we detected, more than half led to premature termination codons, splice acceptor/donor, or frameshift mutations, which would likely lead to the production of aberrant mRNAs subject to nonsense-mediated decay and/or the generation of unstable, truncated PARP1 protein. As we previously noted for premature-termination mutations in yeast *TOP1*, these mutations did not cluster in any particular domains of the *Parp1* open reading frame (Figure 4C). Furthermore, similar to what we observed in yeast, EMS treatment resulted in an overrepresentation of single nucleotide variants, compared to frameshift mutations (Figure 4C).

Strikingly, missense mutations detected in *Parp1* were more frequent in the N-terminal part of the protein, where the DNA-binding domains reside, while no such mutations were observed in the catalytic domain (Figure 4C). Analysis of PARP1 protein levels indicated that other than G400R and A610V, which resulted in complete loss or marked reduction of the PARP1 protein product, no other of the identified missense mutations impacted on PARP protein stability (Figure 5A, Supplementary Figure 3). Computational predictions for the likely consequences of these remaining missense mutations on protein structure and function suggested that all of them were likely to be functionally deleterious (Figure 4D). Due to the fact that all these missense mutations localized within domains known to be involved in DNA binding, we examined their locations relative to the DNA-protein interface as defined by previously published PARP1 structures (26, 44) (Figure 5B). Notably, most of the missense mutations affecting residues in the DNA binding domains clustered at the DNA-protein interface, and did so in proximity to residues that make key DNA contacts (26).

**Figure 4.**
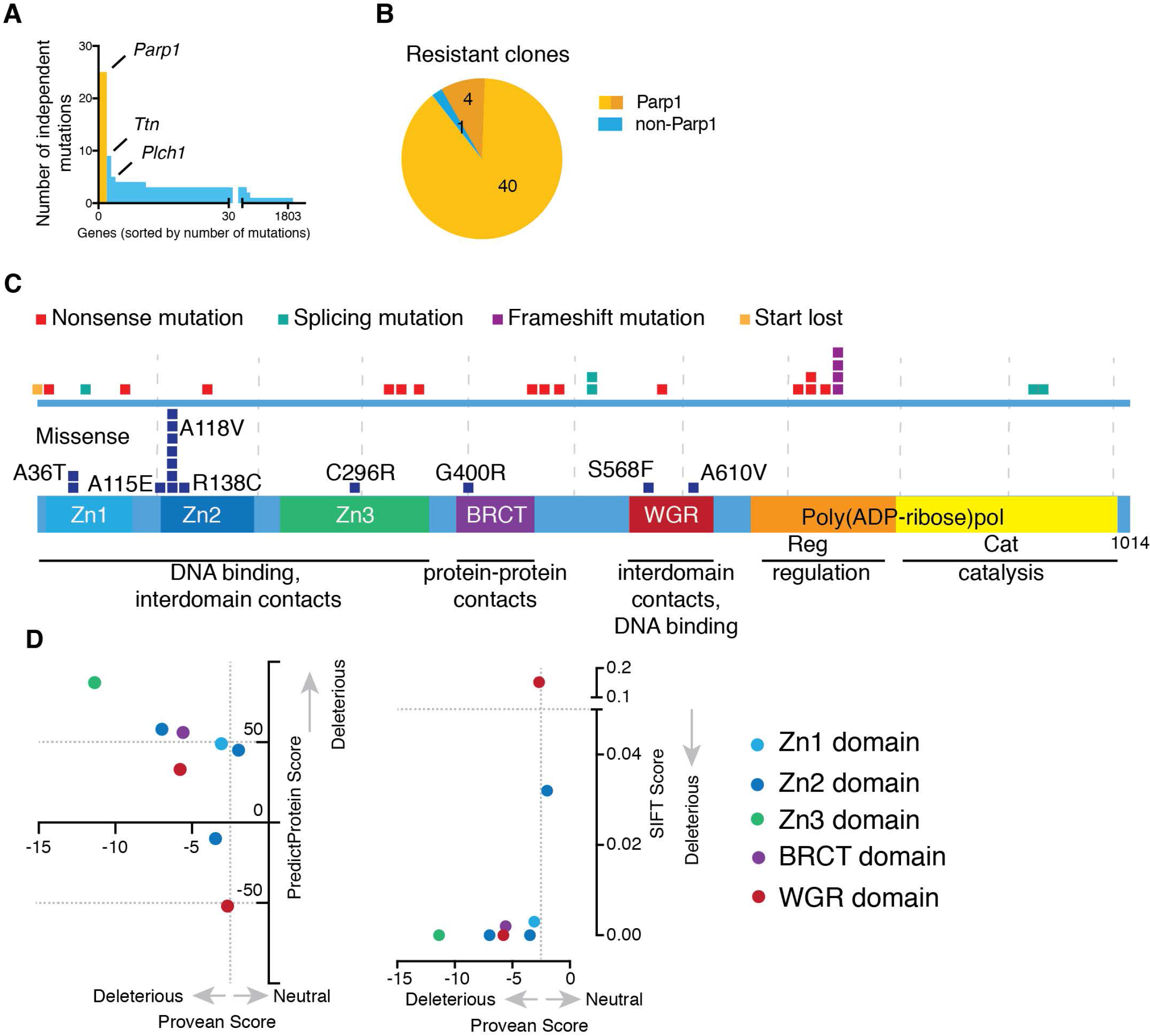
A screen for olaparib resistance indicates critical residues in PARP1. (A) Genes found mutated in the olaparib resistant clones sorted by the number of non-synonymous, independent mutations per gene. (B) Fraction of strains identified by analysis workflow to carry a mutation in *Parp1.* (C) Locations of mutations with respect to the protein sequence. The lower panel shows the distribution of missense mutations, while the upper panel contains all mutations likely to lead to a loss of PARP1 protein: nonsense, frameshift and splice acceptor/donor nucleotide mutations as well as mutations of the start codon. (D) The likelihood of being deleterious of *Parp1* missense mutations as predicted by PredictProtein (y-axis; a score above 50 is considered deleterious) and PROVEAN (x-axis; a score below −2.5 is considered deleterious). Datapoints are coloured by the domain the missense mutations occur in: Light blue, dark blue and green are the three Zinc fingers, respectively, purple denotes the BRCT domain, while red indicates the WGR domain.

Without any a priori knowledge about how olaparib causes cell toxicity, the above data would have suggested that such toxicity is largely driven by a mechanism connected to PARP1 DNA binding. To test this idea, we assessed the missense mutations identified in the DNA binding domains of PARP1 for their potential effects on the ability of PARP1 to bind a double-stranded DNA oligonucleotide. Significantly, this analysis revealed that all the point mutants that did not reduce PARP1 levels showed reduced levels of DNA binding when compared to the wild-type PARP1 protein (Figure 5C, Supplementary Figure 3). Consistent with PARP1 DNA binding triggering its auto-modification by poly ADP-ribose, we found that the PARP1 S568F mutation, which impairs DNA binding, did not exhibit evidence of parylation when cells were treated with hydrogen peroxide (Figure 5D). These findings were therefore in accord with the fact that toxicities of PARP inhibitors such as olaparib are linked to their ability to trap PARP1 on DNA by blocking its catalytic activity (42).

The last clone without an assigned suppressor mutation (C1) may also carry one or more mutations in the non-exonic regions of the *Parp1* gene, or epigenetic modifications altering PARP1 expression, since we could not detect the presence of PARP1 protein in this clone (Figure 5A). Taken together, these results are consistent with a model in which olaparib resistance can arise either from loss of PARP1 or from its decreased ability to bind DNA (Figure 5E).

**Figure 5.**
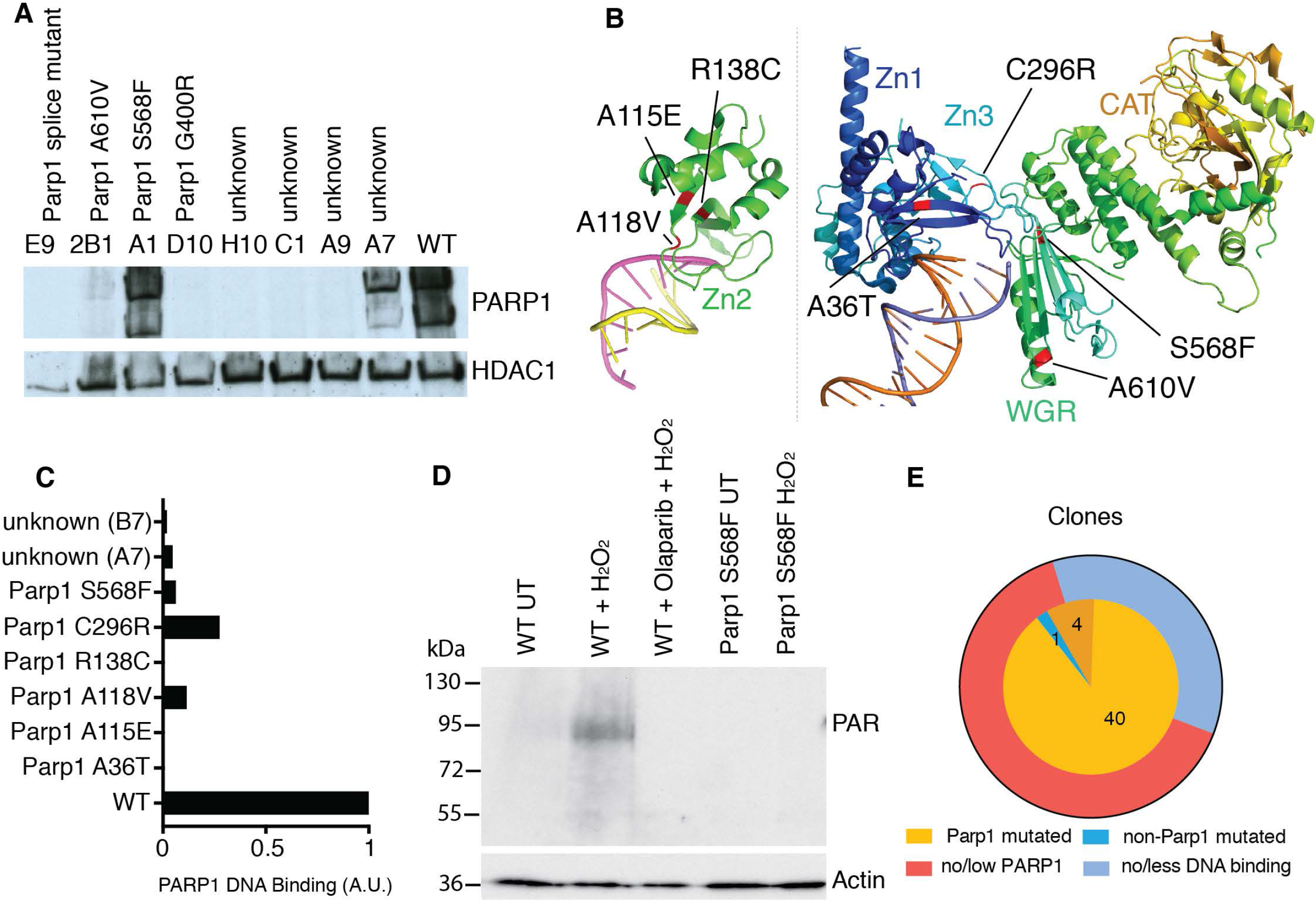
Missense mutations interfere with PARP1 DNA binding. (A) Olaparib resistant clones were assayed for the presence of PARP1 protein. A representative selection is shown; more clones can be seen in Supplementary Figure 3a. (B) Location of missense mutations that do not lead to a loss of PARP1 protein are indicated on partial PARP1 structures of the ZN2 domain by itself (PDB code 3ODC) and the PARP1 protein except ZN2 and the BRCT domain (PDB code 4DQY). (C) Clones carrying mutations that do not lead to loss of/reduction in PARP1 protein were assayed for the protein’s ability to bind a double-stranded DNA oligonucleotide. One experiment each was quantified. All blots can be viewed in Supplementary Figure 3b. (D) Cells with either wild type or mutant PARP1 protein were assayed for their ability to PARylate. Cells were either left untreated or treated with hydrogen peroxide (H2O2). Wild type cells were also treated with olaparib overnight before H2O2 treatment. (E) Summary of olaparib resistant clone analysis. The inner circle shows the mutations that have been identified in every clone (dark yellow signifies mutations identified by manual inspection of the data). The outer circle shows the respective effect on PARP1 protein accounting for the mechanism of olaparib resistance.

## DISCUSSION

Various approaches have been described for systematic identification of genetic and chemo-genetic interactions. Until recently, this search has been largely conducted using approaches based on gene inactivation, either in arrayed or pooled assays. While these approaches have played crucial roles in determining gene-gene and gene-drug interactions, their limited power of resolution does not in general provide information regarding the functional protein domains relevant for the identified interaction. While transposon-based mutagenesis has recently been shown to provide some information at the domain level, this approach is only applicable to loss-of-function mutations, and is biased towards C-terminal domains of proteins (45). In contrast, SNV based approaches can provide a higher level of resolution, and in many cases produce unanticipated results (10, 46–48). Lack of rapid and facile procedures to bridge the phenotype-to-genotype gap has until recently, however, precluded the use of these techniques on a high-throughput scale.

The approach we have described allows the identification of SNV driving drug resistance or resistance to essentially any selective growth condition in a systematic and unbiased way (other than any bias imposed by the mutagenic agent of choice). Importantly, this approach can equally be applied to yeast and to more complex eukaryotes, bringing the power of high-resolution genetic screens to mammalian systems. While we acknowledge that we have carried out our screens with strong selectable cell-viability phenotypes, we envisage applicability in more complex scenarios, for example involving FACS-based selection or cell migration, motility or attachment assays. Highlighting this potential, our results show how, with no previous knowledge, Top1 and PARP1 would have been identified as the most likely targets for the drugs camptothecin and olaparib, respectively.

Toxicity to PARP inhibitors was initially linked to the involvement of PARP1 in the repair of single-strand DNA breaks (49, 50), but more recent data challenged this view (51). The fact that loss of PARP1 drives resistance to PARP inhibitors in wild type genetic backgrounds (52) indeed suggests that inhibition of PARP1 catalytic activity — and not the accumulation of unrepaired DNA lesions — is the major effector of toxicity in such genetic backgrounds. Indeed, recent findings suggest that PARP1 trapping onto DNA, caused by inhibition of its catalytic activity, is the main cause of toxicity (42). Our data further support this model, as all the suppressor clone variants which we identified that did not result in loss of PARP1 protein negatively affected its binding to DNA. Preventing PARP1 binding to DNA thus appears to be sufficient to circumvent the toxicity of PARP inhibitors, and the fact that we identified no mutants specifically defective in catalytic function reinforces the idea of trapped PARP1 as the main cytotoxic lesion for olaparib in wild-type mammalian cells. This may not only be important for our understanding how PARP inhibitors function but also for mechanisms of intrinsic or evolved tumor resistance towards such clinical agents in patients.

As we have exemplified by our analyses of TOP1 and PARP1, the level of detail on critical functional domains and residues increases with the number of samples sequenced. Because of their genome size, screens based in mammalian systems require greater sequencing power than screens conducted in simpler organisms such as yeast. Moreover, as compared to yeasts, the more complex genome architecture in mammalian systems - where there is more intergenic DNA, a larger number of genes and an abundance of intronic sequences - increases the chances of isolating variants affecting protein levels, rather than protein function. One solution to bypass such issues will be to run two-tiered screens, initially using whole exome sequencing on a subset of suppressors in order to identify top gene hits driving resistance, and then using targeted exome sequencing to test the rest of the samples, either through analysis of various individual clones or bulk sequencing of the resistant population. In addition, we can envision alternative scenarios where a gene identified in an initial screen could be marked/tagged in a way to allow selection of mutations that affect protein function but not protein levels. This approach can also be combined with CRISPR-Cas9-mediated *in vivo* targeted mutagenesis, via a library of gRNAs directed towards the exonic regions of the gene (Christopher Lord, personal communication). We anticipate that such developments, along with expected further increases in sequencing throughput and associated cost reductions, will pave the way for hitherto unprecedented genetic analyses on comprehensive and systematic scales.

## Data Availability

Yeast DNA sequencing data are available from European Nucleotide Archive (ENA) under the accession code PRJEB2977, and mouse DNA sequencing data are available under accession code PRJEB13612. Access codes for specific samples are detailed in Supplementary Table 2. Source data for Figures 2 and 3 are provided in Supplementary Table 2; source data for Figures 4 and 5 are provided in Supplementary Table 3 and Supplementary Figure 3.

## Code Availability

Custom code used to analyse sequencing data and to draw figures is available in the following repository: http://github.com/fabiopuddu/Herzog2018

## Acknowledgements

We thank Carmen Diaz Soria for help with the yeast screens; Josef Penninger for the gift of the AN3-12 mESC line; Carla Daniela Robles Espinoza and Martin del Castillo Velasco-Herrera for critical reading of the manuscript; James Hewinson for assistance with the submission for sequencing samples; and all the members of the SPJ laboratory for helpful discussions. The project was conceived by JVF, FP, MH and SPJ. Yeast mutagenesis, suppressor screening, and DNA extractions were carried out by FP and NJG. Mouse haploid cell purification, EMS mutagenesis, and screening were carried out by JVF; DNA extractions and PARP assays by JC. Analysis of yeast and mouse whole genome/exome sequencing data was carried out by FP and MH, respectively. The manuscript was largely written by MH, FP, and SPJ, with contributions made by all the other authors.

---

**Supplementary Figure 1.**
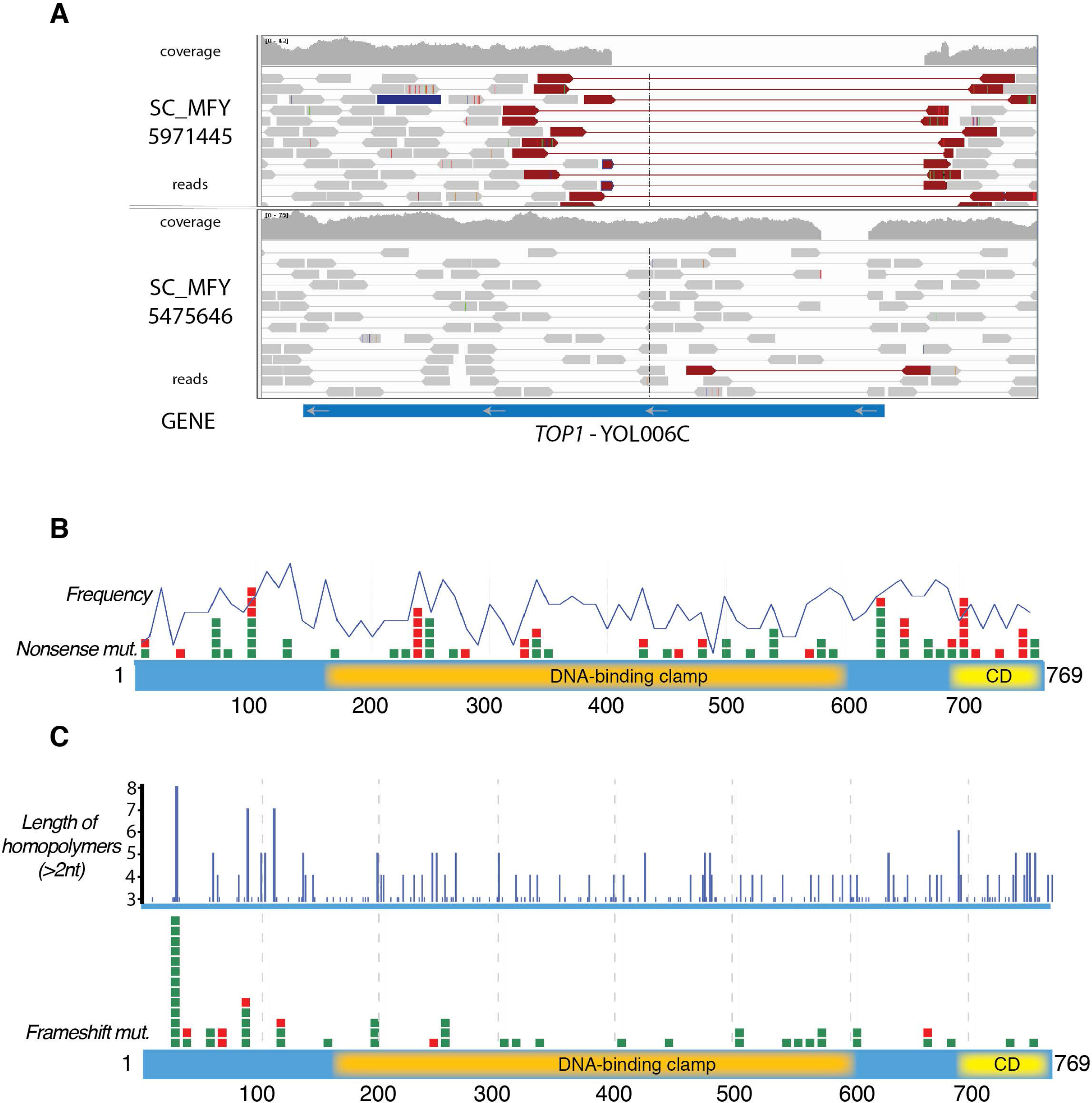
(A) Examples of two camptothecin resistant yeast strains, which each carry a large deletion in the *TOP1* gene. (B) Nonsense mutations are depicted as in Figure 2D. Superimposed is the frequency of codons that can be mutated to a stop codon by one nucleotide change. (C) Frameshift mutations are depicted as in Figure 2E. Above the locations of homopolymers of a length of at least 3nt in the *TOP1* gene are plotted, their length indicated on the y-axis.

**Supplementary Figure 2.**
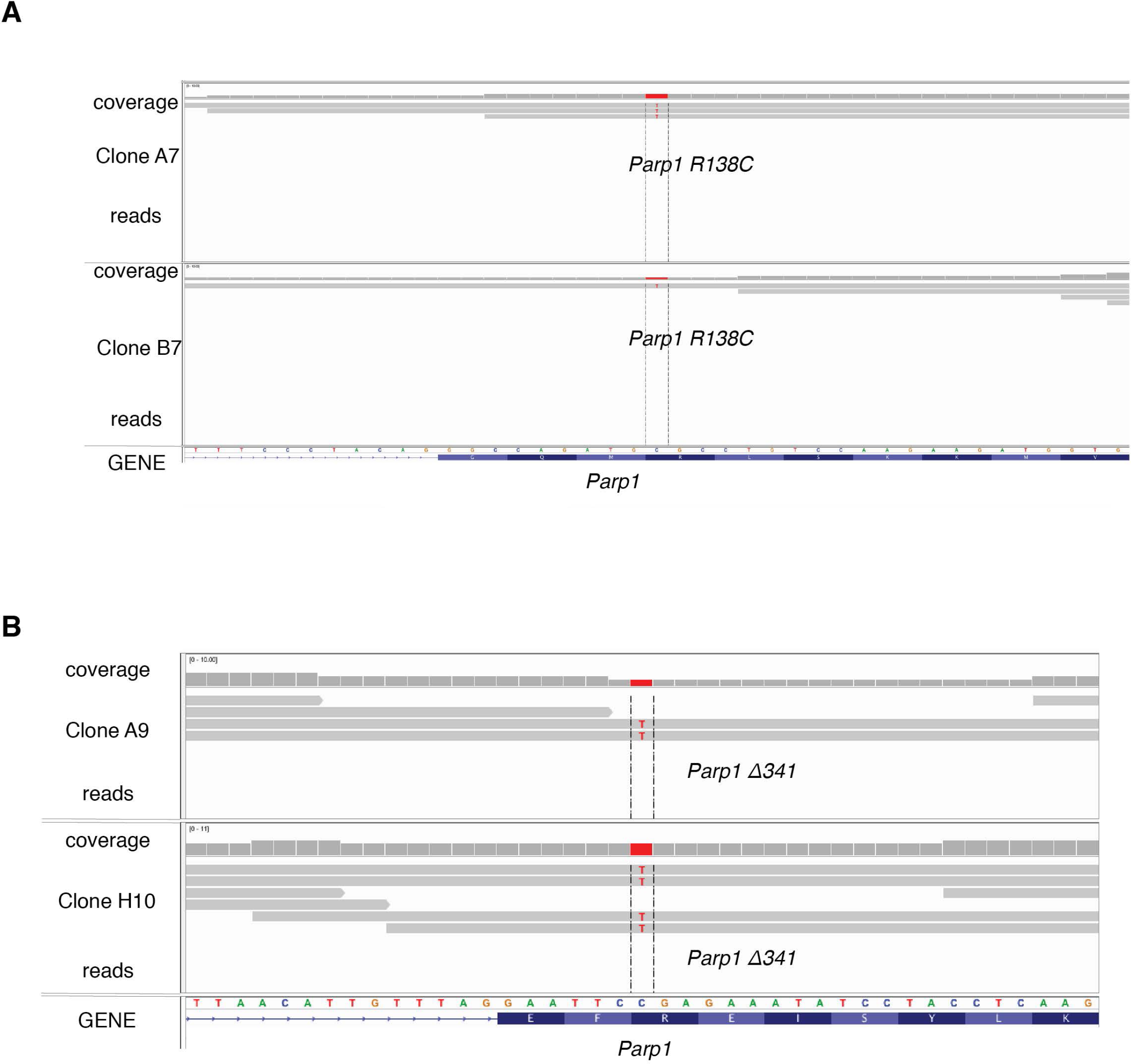
(A) Integrative Genomics Viewer (IGV) panels of the sequencing data for clones A9 and H10 showing the *Parpl* mutation *(Parpl Δ341)* that did not pass the filters. (B) Integrative Genomics Viewer (IGV) panels of the sequencing data for clones A7 and B7 showing the *Parp1* mutation *(Parp1 R138C)* that did not pass the filters.

**Supplementary Figure 3.**
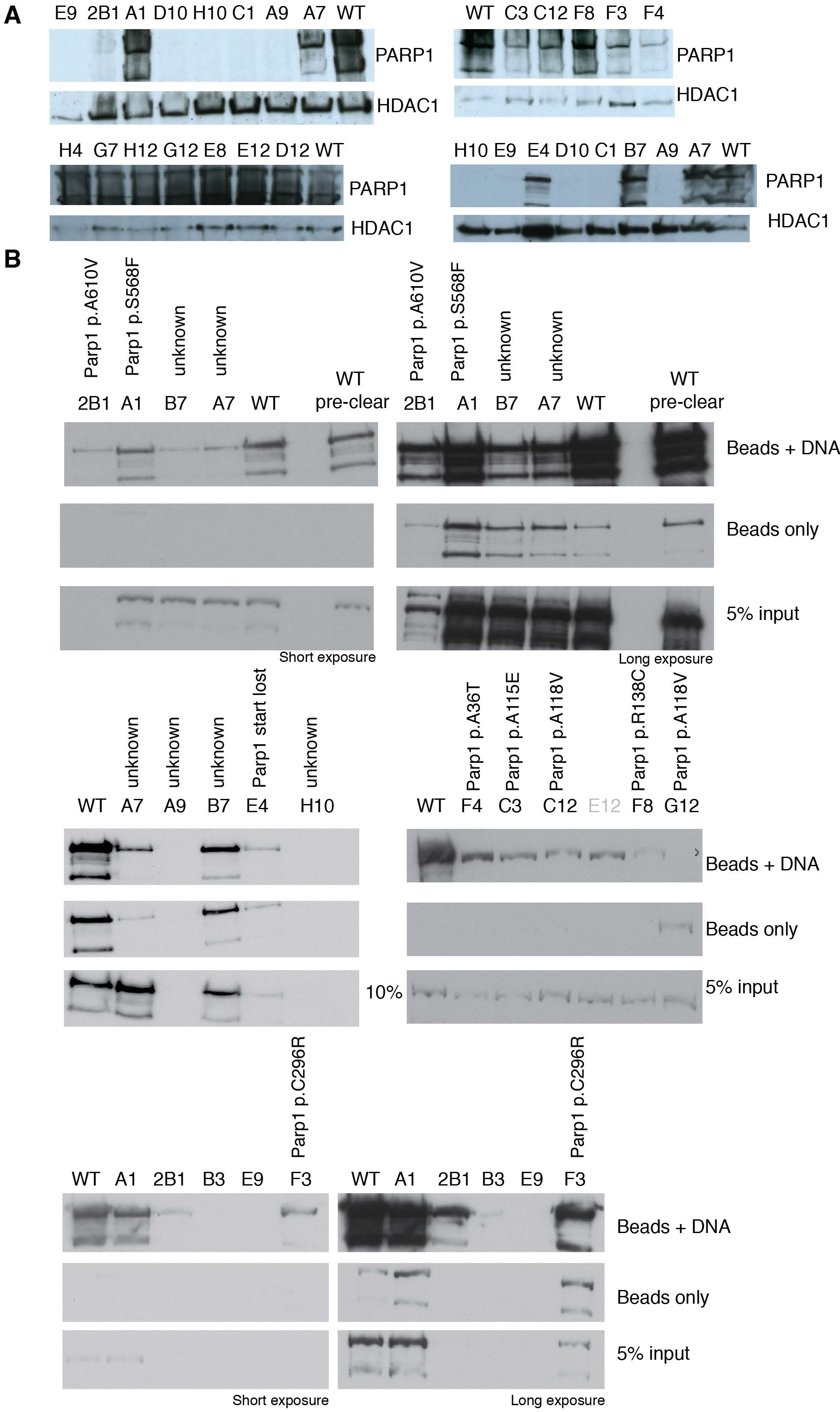
(A) Western blots for PARP1 protein for olaparib resistant clones. (B) PARP1-DNA binding assays in olaparib resistant clones. Two exposure each are shown.

## REFERENCES

1. Hillenmeyer,M.E., Fung,E., Wildenhain,J., Pierce,S.E., Hoon,S., Lee,W., Proctor,M., St Onge,R.P., Tyers,M., Koller,D., et al. (2008) The chemical genomic portrait of yeast: uncovering a phenotype for all genes. Science (New York, N.Y.), 320, 362–365.

2. Parsons,A.B., Lopez,A., Givoni,I.E., Williams,D.E., Gray,C.A., Porter,J., Chua,G., Sopko,R., Brost,R.L., Ho,C.-H., et al. (2006) Exploring the mode-of-action of bioactive compounds by chemical-genetic profiling in yeast. Cell, 126, 611–625.

3. Nijman,S.M.B. (2015) Functional genomics to uncover drug mechanism of action. Nat. Chem. Biol., 11, 942–948.

4. van Leeuwen,J., Pons,C., Mellor,J.C., Yamaguchi,T.N., Friesen,H., Koschwanez,J., Ušaj,M.M., Pechlaner,M., Takar,M., Ušaj,M., et al. (2016) Exploring genetic suppression interactions on a global scale. Science (New York, N.Y.), 354, aag0839–aag0839.

5. Forsburg,S.L. (2001) The art and design of genetic screens: yeast. Nature reviews. Genetics, 2, 659–668.

6. Grimm,S. (2004) The art and design of genetic screens: mammalian culture cells. Nature reviews. Genetics, 5, 179–189.

7. Horner,V.L. and Caspary,T. (2011) Creating a ‘hopeful monster’: mouse forward genetic screens. Methods Mol. Biol., 770, 313–336.

8. Trahey,M. and McCormick,F. (1987) A cytoplasmic protein stimulates normal N-ras p21 GTPase, but does not affect oncogenic mutants. Science (New York, N.Y.), 238, 542–545.

9. Ingram,V.M. (1957) Gene mutations in human haemoglobin: the chemical difference between normal and sickle cell haemoglobin. Nature, 180, 326–328.

10. Rolef Ben-Shahar,T., Heeger,S., Lehane,C., East,P., Flynn,H., Skehel,M. and Uhlmann,F. (2008) Eco1-dependent cohesin acetylation during establishment of sister chromatid cohesion. Science (New York, N.Y.), 321, 563–566.

11. Carette,J.E., Guimaraes,C.P., Varadarajan,M., Park,A.S., Wuethrich,I., Godarova,A., Kotecki,M., Cochran,B.H., Spooner,E., Ploegh,H.L., et al. (2009) Haploid genetic screens in human cells identify host factors used by pathogens. Science (New York, N.Y.), 326, 1231–1235.

12. Leeb,M. and Wutz,A. (2011) Derivation of haploid embryonic stem cells from mouse embryos. Nature, 479, 131–134.

13. Elling,U., Taubenschmid,J., Wirnsberger,G., O'Malley,R., Demers,S.-P., Vanhaelen,Q., Shukalyuk,A.I., Schmauss,G., Schramek,D., Schnuetgen,F., et al. (2011) Forward and reverse genetics through derivation of haploid mouse embryonic stem cells. Cell Stem Cell, 9, 563–574.

14. Munroe,R. and Schimenti,J. (2009) Mutagenesis of mouse embryonic stem cells with ethylmethanesulfonate. Methods Mol. Biol., 530, 131–138.

15. Li,H. and Durbin,R. (2009) Fast and accurate short read alignment with Burrows-Wheeler transform. Bioinformatics (Oxford, England), 25, 1754–1760.

16. Li,H., Handsaker,B., Wysoker,A., Fennell,T., Ruan,J., Homer,N., Marth,G., Abecasis,G. and Durbin,R. (2009) The Sequence Alignment/Map format and SAMtools. Bioinformatics (Oxford, England), 25, 2078–2079.

17. McLaren,W., Pritchard,B., Rios,D., Chen,Y., Flicek,P. and Cunningham,F. (2010) Deriving the consequences of genomic variants with the Ensembl API and SNP Effect Predictor. Bioinformatics (Oxford, England), 26, 2069–2070.

18. Danecek,P., Auton,A., Abecasis,G., Albers,C.A., Banks,E., DePristo,M.A., Handsaker,R.E., Lunter,G., Marth,G.T., Sherry,S.T., et al. (2011) The variant call format and VCFtools. Bioinformatics (Oxford, England), 27, 2156–2158.

19. Keane,T.M., Goodstadt,L., Danecek,P., White,M.A., Wong,K., Yalcin,B., Heger,A., Agam,A., Slater,G., Goodson,M., et al. (2011) Mouse genomic variation and its effect on phenotypes and gene regulation. Nature, 477, 289–294.

20. Narzisi,G., O'Rawe,J.A., Iossifov,I., Fang,H., Lee,Y.-H., Wang,Z., Wu,Y., Lyon,G.J., Wigler,M. and Schatz,M.C. (2014) Accurate de novo and transmitted indel detection in exome-capture data using microassembly. Nature methods, 11, 1033–1036.

21. Ng,P.C. and Henikoff,S. (2006) Predicting the effects of amino acid substitutions on protein function. Annu Rev Genomics Hum Genet, 7, 61–80.

22. Ng,P.C. and Henikoff,S. (2001) Predicting deleterious amino acid substitutions. Genome research, 11, 863–874.

23. Ng,P.C. and Henikoff,S. (2003) SIFT: Predicting amino acid changes that affect protein function. Nucleic acids research, 31, 3812–3814.

24. Kumar,P., Henikoff,S. and Ng,P.C. (2009) Predicting the effects of coding non-synonymous variants on protein function using the SIFT algorithm. Nature protocols, 4, 1073–1081.

25. Yachdav,G., Kloppmann,E., Kajan,L., Hecht,M., Goldberg,T., Hamp,T., Honigschmid,P., Schafferhans,A., Roos,M., Bernhofer,M., et al. (2014) PredictProtein--an open resource for online prediction of protein structural and functional features. Nucleic acids research, 42, W337–43.

26. Langelier,M.-F., Planck,J.L., Roy,S. and Pascal,J.M. (2012) Structural basis for DNA damage-dependent poly(ADP-ribosyl)ation by human PARP-1. Science (New York, N.Y.), 336, 728–732.

27. Hsiang,Y.H., Hertzberg,R., Hecht,S. and Liu,L.F. (1985) Camptothecin induces protein-linked DNA breaks via mammalian DNA topoisomerase I. The Journal of biological chemistry, 260, 14873–14878.

28. Bjornsti,M.-A., Benedetti,P., Viglianti,G.A. and Wang,J.C. (1989) Expression of human DNA topoisomerase I in yeast cells lacking yeast DNA topoisomerase I: restoration of sensitivity of the cells to the antitumor drug camptothecin. Cancer Res., 49, 6318–6323.

29. Pommier,Y., Sun,Y., Huang,S.-Y.N. and Nitiss,J.L. (2016) Roles of eukaryotic topoisomerases in transcription, replication and genomic stability. Nature reviews. Molecular cell biology, 17, 703–721.

30. Rouse,J. (2004) Esc4p, a new target of Mec1p (ATR), promotes resumption of DNA synthesis after DNA damage. The EMBO journal, 23, 1188–1197.

31. Puddu,F., Oelschlaegel,T., Guerini,I., Niu,H., Ochoa-Montaño,B., Viré,E., Sung,P., Keane,T.M., Adams,D.J., Keane,T.M., et al. (2015) Synthetic viability genomic screening defines Sae2 function in DNA repair. The EMBO journal, 34, 1509–1522.

32. Huertas,P., Cortés-Ledesma,F., Sartori,A.A., Aguilera,A. and Jackson,S.P. (2008) CDK targets Sae2 to control DNA-end resection and homologous recombination. Nature, 455, 689–692.

33. Mimitou,E.P. and Symington,L.S. (2010) Ku prevents Exo1 and Sgs1-dependent resection of DNA ends in the absence of a functional MRX complex or Sae2. EMBO J, 29, 3358–3369.

34. Coulondre,C. and Miller,J.H. (1977) Genetic studies of the lac repressor. IV. Mutagenic specificity in the lacI gene of Escherichia coli. J. Mol. Biol., 117, 577–606.

35. Puddu,F., Salguero,I., Herzog,M., Geisler,N.J., Costanzo,V. and Jackson,S.P. (2017) Chromatin determinants impart camptothecin sensitivity. EMBO reports, 10.15252/embr.201643560.

36. Mitchell,P. and Tollervey,D. (2003) An NMD pathway in yeast involving accelerated deadenylation and exosome-mediated 3‘-->5’ degradation. Molecular Cell, 11, 1405–1413.

37. Lynn,R.M., Bjornsti,M.-A., Caron,P.R. and Wang,J.C. (1989) Peptide sequencing and site-directed mutagenesis identify tyrosine-727 as the active site tyrosine of Saccharomyces cerevisiae DNA topoisomerase I. Proc Natl Acad Sci U S A, 86, 3559–3563.

38. Eng,W.K., Pandit,S.D. and Sternglanz,R. (1989) Mapping of the active site tyrosine of eukaryotic DNA topoisomerase I. The Journal of biological chemistry, 264, 13373–13376.

39. Redinbo,M.R., Stewart,L., Kuhn,P., Champoux,J.J. and Hol,W.G. (1998) Crystal structures of human topoisomerase I in covalent and noncovalent complexes with DNA. Science (New York, N.Y.), 279, 1504–1513.

40. Stewart,L., Redinbo,M.R., Qiu,X., Hol,W.G. and Champoux,J.J. (1998) A model for the mechanism of human topoisomerase I. Science (New York, N.Y.), 279, 1534–1541.

41. Forment,J.V., Herzog,M., Coates,J., Konopka,T., Gapp,B.V., Nijman,S.M., Adams,D.J., Keane,T.M. and Jackson,S.P. (2017) Genome-wide genetic screening with chemically mutagenized haploid embryonic stem cells. Nat. Chem. Biol., 13, 12–14.

42. Lord,C.J. and Ashworth,A. (2017) PARP inhibitors: Synthetic lethality in the clinic. Science (New York, N.Y.), 355, 1152–1158.

43. Murai,J., Huang,S.-Y.N., Das,B.B., Renaud,A., Zhang,Y., Doroshow,J.H., Ji,J., Takeda,S. and Pommier,Y. (2012) Trapping of PARP1 and PARP2 by Clinical PARP Inhibitors. Cancer Res., 72, 5588–5599.

44. Langelier,M.-F., Planck,J.L., Roy,S. and Pascal,J.M. (2011) Crystal structures of poly(ADP-ribose) polymerase-1 (PARP-1) zinc fingers bound to DNA: structural and functional insights into DNA-dependent PARP-1 activity. The Journal of biological chemistry, 286, 10690–10701.

45. Michel,A.H., Hatakeyama,R., Kimmig,P., Arter,M., Peter,M., Matos,J., De Virgilio,C. and Kornmann,B. (2017) Functional mapping of yeast genomes by saturated transposition. eLife, 6, E3179.

46. Hardy,C.F., Dryga,O., Seematter,S., Pahl,P.M. and Sclafani,R.A. (1997) mcm5/cdc46-bob1 bypasses the requirement for the S phase activator Cdc7p. Proc Natl Acad Sci U S A, 94, 3151–3155.

47. Sandrock,T.M., O'Dell,J.L. and Adams,A.E. (1997) Allele-specific suppression by formation of new protein-protein interactions in yeast. Genetics, 147, 1635–1642.

48. Zhao,X., Chabes,A., Domkin,V., Thelander,L. and Rothstein,R. (2001) The ribonucleotide reductase inhibitor Sml1 is a new target of the Mec1/Rad53 kinase cascade during growth and in response to DNA damage. The EMBO journal, 20, 3544–3553.

49. Farmer,H., McCabe,N., Lord,C.J., Tutt,A.N.J., Johnson,D.A., Richardson,T.B., Santarosa,M., Dillon,K.J., Hickson,I., Knights,C., et al. (2005) Targeting the DNA repair defect in BRCA mutant cells as a therapeutic strategy. Nature, 434, 917–921.

50. Bryant,H.E., Schultz,N., Thomas,H.D., Parker,K.M., Flower,D., Lopez,E., Kyle,S., Meuth,M., Curtin,N.J. and Helleday,T. (2005) Specific killing of BRCA2-deficient tumours with inhibitors of poly(ADP-ribose) polymerase. Nature, 434, 913–917.

51. Helleday,T. (2011) The underlying mechanism for the PARP and BRCA synthetic lethality: clearing up the misunderstandings. Mol Oncol, 5, 387–393.

52. Pettitt,S.J., Rehman,F.L., Bajrami,I., Brough,R., Wallberg,F., Kozarewa,I., Fenwick,K., Assiotis,I., Chen,L., Campbell,J., et al. (2013) A genetic screen using the PiggyBac transposon in haploid cells identifies Parp1 as a mediator of olaparib toxicity. PloS one, 8, e61520.

